# Estimation of the Achilles tendon twist in vivo by individual triceps surae muscle stimulation

**DOI:** 10.1101/2024.02.28.582458

**Authors:** Lecompte Laura, Crouzier Marion, Baudry Stéphane, Vanwanseele Benedicte

## Abstract

The Achilles tendon (AT) is comprised of three distinct subtendons, each arising from the one of the three heads of the triceps surae muscles: gastrocnemius medialis (GM), gastrocnemius lateralis (GL) and soleus (SOL). These subtendons exhibit a twisted structure, classified as low (Type I), medium (Type II), and high (Type III) twist, based on cadaveric studies. Nevertheless, the in-vivo investigation of AT twist is notably scarce, resulting in a limited understanding of its functional significance. The aim of this study was to give insights into the complex 3D AT structure in vivo. 30 healthy participants underwent individual stimulation of each of the triceps surae muscles at rest with the foot attached to the pedal of an isokinetic dynamometer. Ultrasound images were captured to concomitantly examine the displacement of the superficial, middle and deep AT layers. SOL stimulation resulted in the highest AT displacement followed by GM and GL stimulation. Independent of the muscle stimulated, non-uniformity within the AT was observed with the deep layer exhibiting more displacement compared to the middle and superficial layers, hence important inter-individual differences in AT displacement were noticeable. By leveraging these individual displacement patterns during targeted stimulations in conjunction with cadaveric twist classifications providing insights into the area of each specific subtendon, our classification identified 19 subjects with a ’low’ and 11 subjects with a ’high’ AT twist. More research is needed to understand the complexity of the AT twisted structure in vivo to further understand its effect on AT properties and behaviour.

## 1. INTRODUCTION

The Achilles tendon (AT) plays a crucial role in human locomotion and reveals a distinctive yet complex anatomy, characterized by the presence of three subtendons originating from the gastrocnemius medialis (GM), gastrocnemius lateralis (GL) and soleus (SOL) muscles. The main function of the AT is to transmit the forces generated by these muscles to the calcaneus. By acting as an energy-storing tendon through a process of stretching and recoiling, the AT can further reduce the energetic cost associated with locomotion. What truly distinguishes the AT from other tendons in the human body is its descending twisted structure, initially described in human by Cummins and Anson (1946). The AT twist is characterized by a proximal to distal clockwise rotation in the left limb and a counter-clockwise rotation in the right limb. This unique twist has been categorized into three types of rotation: type I, with the least amount of twist, type II, characterized by moderate level of twisting, and type III, showing the highest degree of twist. Several cadaveric studies have reported the prevalence of this various twist pattern, although in different populations (Szaro et al. 2009, Edama et al. 2015, Pekala et al. 2017). The twist classification agrees between studies although the categories might differ slightly. For example, specimens classified as Type I and II by Pekala et al. (2017) would correspond to specimens classified Type II and III by Edama et al (2015). Accordingly, Type I is the most common in Pekala and colleague’s findings, while it is Type II in the study by Edama and colleagues. Table 1 summarizes the proportions of Achilles tendon twist types in different studies highlighting a consensus in the literature that low twist has the highest prevalence among humans. Despite the growing interest, there is currently only one study that estimated this twist in vivo using high field 7-Tesla MRI (Cone et al. 2023) resulting in a very limited understanding of it’s in vivo structure.

**Table 1:** Overall distribution of Achilles tendon twist types in different studies.

| Study | N | Type I | Type II | Type III |
| --- | --- | --- | --- | --- |
| **Cummins and Anson (1946)** | 100 | 52% | 35% | 13% |
| **Edama et al. (2015)** | 110 | 50% | 43% | 7% |
| **Edama et al. (2016)** | 132 | 24% | 67% | 9% |
| **Pekala et al. (2017)** | 106 | 48% | 46% | 6% |
| **Edama et al. (2021)** | 30 | 13% | 77% | 10% |
| **Cone et al. (2023)** | 20 | 80% | 12% | 8% |
| **Pringels et al. (2023)** | 18 | 50% | 50% | 0% |

**Table 2:** Overview of the parameters collected during stimulation of GM, GL and SOL in neutral foot position in all subjects (n = 30).

|  | GL stimulation | GM stimulation | SOL stimulation |
| --- | --- | --- | --- |
| *Stimulation intensity (mA)* |  |  |  |
| | $25.8 \pm 4.5$ | $26.3 \pm 4.2$ | $32.0 \pm 4.4$ |
| *Torque (Nm)* |  |  |  |
| | $9.4 \pm 3.5$ | $11.1 \pm 3.2$ | $12.5 \pm 5.8$ |
| *Amplitude M-wave (mV)* |  |  |  |
| | $2.7 \pm 2.1$ | $2.6 \pm 1.5$ | $3.2 \pm 2.6$ |
| *AT absolute displacement (mm)* |  |  |  |
| Deep Layer | $0.84 \pm 0.49$ | $1.45 \pm 0.91$ | $1.58 \pm 1.17$ |
| Mid Layer | $0.61 \pm 0.42$ | $1.12 \pm 0.82$ | $1.26 \pm 1.05$ |
| Superficial layer | $0.53 \pm 0.37$ | $0.99 \pm 0.76$ | $1.09 \pm 0.92$ |
| Mean of the three layers | $0.66 \pm 0.42$ | $1.18 \pm 0.82$ | $1.31 \pm 1.04$ |
| *Absolute AT displacement normalized (% of sum displacement all layers)* |  |  |  |
| Deep Layer | $43.9 \pm 5.2$ | $41.6 \pm 4.8$ | $42.1 \pm 4.6$ |
| Mid Layer | $30.8 \pm 4.2$ | $30.9 \pm 3.4$ | $31.1 \pm 2.9$ |
| Superficial layer | $25.4 \pm 5.1$ | $27.5 \pm 3.4$ | $26.8 \pm 3.2$ |
| *Intra-tendinous sliding (mm)* |  |  |  |
| | $0.32 \pm 0.12$ | $0.46 \pm 0.24$ | $0.49 \pm 0.36$ |

**Table 3:**
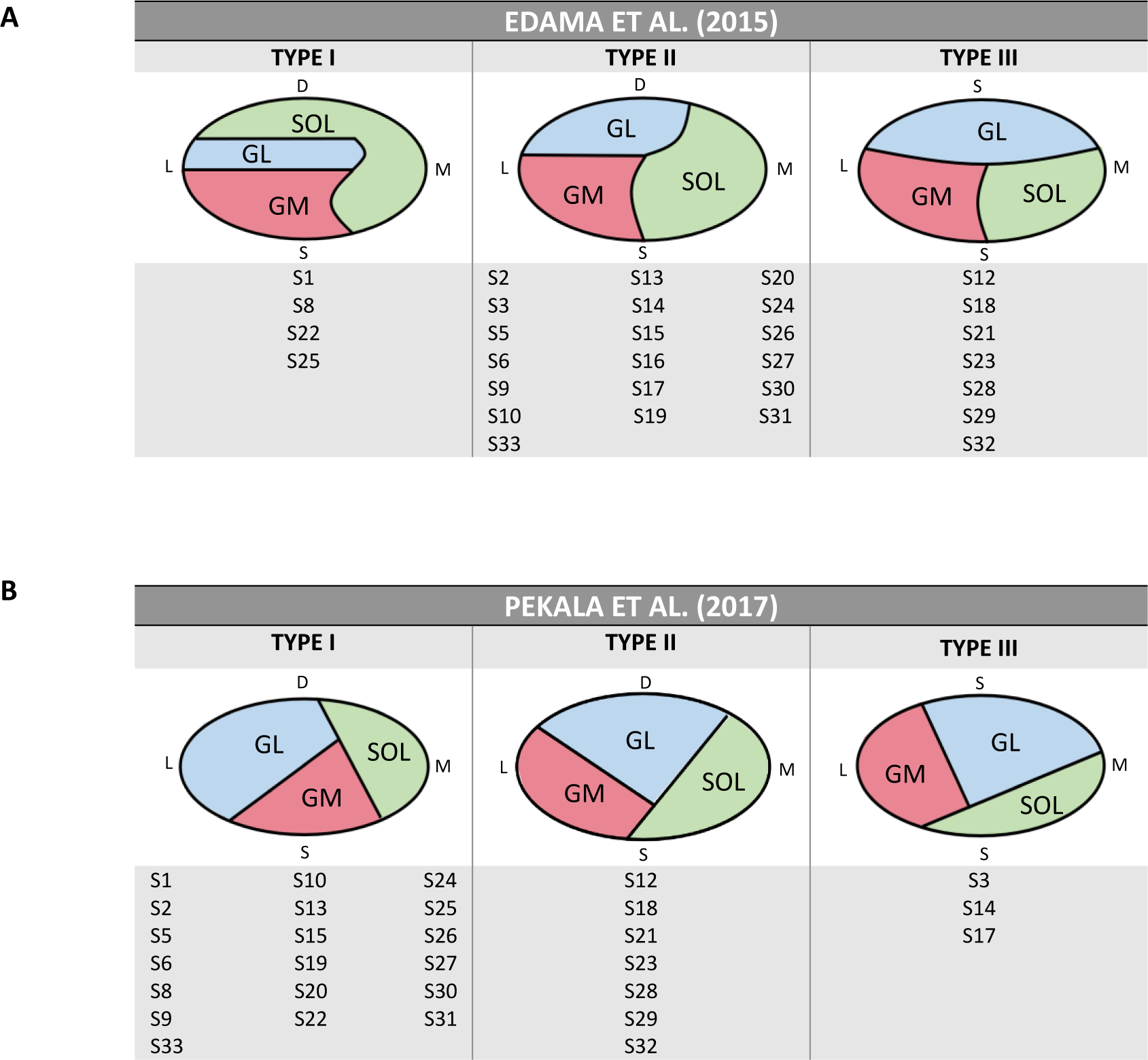
A) Achilles tendon twist distribution based on the twist definitions of Edama et al. (2015). B) Achilles tendon twist distribution following the twist definitions of Pekala et al. (2017). For each twist classification, a schematic representation of all subtendons (GM = gastrocnemius medialis, GL = gastrocnemius lateralis, SOL = soleus) at the insertion into the calcaneal bone is shown: D represents the deep part, S represents the superficial part, L represents the lateral part and M represents the medial part of the Achilles tendon.

A possibility to give insights into the AT structure in vivo is to combine methods that distinguish the displacement of the different Achilles tendon subparts during individual force production from the triceps surae muscles. Given that the myotendinous junction of each triceps surae muscle and so the start of each subtendon is easy to image, coupling this with the capability to distinguish between different Achilles tendon subparts more distally can offer valuable insights into the AT twisting structure.. The use of transcutaneous electrical stimulation allows to target the activation of only one of the triceps surae muscles. Stimulating a muscle individually would thus result in a force transmission that happens mainly within the area of tendon fascicles originating from the stimulated muscle. Since subtendons are not fully independent (Maas et al. 2020), it is expected that the mechanical connections between subtendons will generate motions in other subparts during individual stimulation, but to a reduced extent. This phenomenon occurs due to the sliding interaction between subtendons, which is referred to as intra-tendinous sliding. To the best of our knowledge, only three studies have used myo-electrical stimulation in human and analyzed AT displacements simultaneously (Lehr et al. 2021, Khair et al. 2022, Klaiber et al. 2023). However, none of these studies used such data to infer the amount of AT twist, and none of them stimulated all triceps surae muscles (only two muscles). Coupling the techniques of individual muscle stimulation of all triceps surae muscles and quantification of longitudinal motions occurring in response in the AT sublayers by ultrasound imaging hence appears as a powerful setup to gain better insights in the AT substructure. To complete this investigation, one may want to investigate if the horizontal foot position would impact the intra-tendinous sliding. Voluntary plantarflexions with the foot rotated outwards are known to impact the intra-tendinous sliding, when compared to plantarflexion in a neutral foot position (Crouzier et al. 2022). Yet, it is not known whether this is purely mechanical, i.e. due to untightening the AT by rotating against the twist, or whether this is influenced by other factors such as a different neuromuscular control between foot positions. The use of individual electrical stimulation combined with electromyographic measures of the triceps surae muscles would help unravel such a question.

The aims of this research were to investigate the effect of individual stimulation of GL, GM and SOL (1) on the displacement within the AT and (2) with different horizontal foot orientations. Additionally, we aimed to (3) estimate the AT twist in vivo by examining displacement patterns in the sagittal view of the AT from ultrasound images during selective muscle activation of these triceps surae muscles and associate these findings to the current cadaver twist definitions. We hypothesized that SOL would elicit highest displacement values, being the biggest plantar flexor muscle of the triceps surae muscles, and that toes-out foot position would result in higher intra-tendinous sliding. Further, we hypothesized that we would be able to differentiate individuals into groups based on a low or high twist by linking the individual displacement patterns during selective stimulation to the cadaver twist type definitions.

## 2. METHODS

### 2.1. Participants

For this study, 33 healthy individuals were recruited according to a convenience sampling method Given the insufficient data available in the literature, we did not perform an a-priori sample size calculation. Instead, sample size on prevalence values obtained from cadaver studies, highlighting the low prevalence of Type III twist (ranging from 0 to 13%). To ensure robust representation of all twisting types, we aimed to include a minimum of 30 participants. All participants had no lower limb injuries or orthopedic problems within the preceding 6 months. Prior to the experiment, written informed consent was obtained from all participants and ethical approval for this study was approved by SMEC KU Leuven (number G-2022-6081). 1 participant had to be excluded due to being too sensitive to the muscle stimulations and 2 participants were excluded after our quality control process concerning the M-wave EMG data explained further (*see 2.3.4. EMG*). Thus, we report data for 30 individuals (19 women and 11 men; age: 25.6 ± 2.9 years, BMI: 22.0 ± 2.3 kg/m^2^).

### 2.2. Experimental design

Participants attended one laboratory session in the Movement & Posture Analysis Lab Leuven (Leuven, Belgium). They were lying prone with the knee at 0° (fully extended) and the ankle at 5° plantarflexion. The foot was securely attached to a dynamometer (Biodex Medical Systems, Shirley, New York, USA) and both the thighs and shoulders were strapped to the table.

The protocol started with a standardized warm-up where participants had to execute isometric plantarflexion contractions, gradually increasing from low intensity and progressively reaching approximately 70 to 80% of their self-estimated maximum capacity. To facilitate this, real-time visual feedback was presented on a screen on the side of the participant. Then, participants performed two isometric maximal voluntary contractions (MVC; three if the difference in the maximal force between the first two contractions was more than 10%) for 5s each, with 120s rest in between.

The skin was shaved and cleaned with alcohol, and muscle borders were drawn on the skin with the help of ultrasound imaging to guide the placement of electrodes. First, a pair of 10 mm diameter stimulating electrodes (Ag/AgCl, Technomed, Hertforshire, UK) were placed on each of the triceps surae muscles with an average distance of ∼ 2 cm between both electrodes. Overall, these stimulating electrodes were placed on the prominent bulge of both GM and GL and distally in the middle lower leg but proximal to the free AT for SOL. However, individual adjustments were made to optimize the response for each participant. This involved determining an optimal stimulation location by repositioning the electrodes and visually checking the muscle response following a single 200 µs electrical pulse delivered by a constant current electrical stimulator (DS7R, Digitimer, Hertfordshire, UK). The location with the highest visible muscle contraction was indicated by two marks on the skin (one for each electrode). After the placement of the EMG electrodes, the optimal site of stimulation was checked by recording the muscle compound action potential (M-wave) and moving the stimulating electrodes to obtain the greatest M-wave amplitude for a given stimulation intensity.

A second pair of EMG electrodes (Ag/AgCL electrodes, Ambu A/S, Ballerup, Denmark) with a 20mm interelectrode distance was placed on the muscle belly, according to the SENIAM recommendations as much as possible, even if priority was given to the location of the stimulating electrodes in case of too close locations. EMG signals were sampled at 2000 Hz using an EMG amplifier (ZeroWire EMG, Cometa, Milano, Italy) and displayed on a screen in front of the investigator to check the EMG response in real-time.

Finally, stimulation intensity was determined as the one generating the maximal M-wave on the targeted muscle, without having any contribution from the two surrounding muscles (i.e. no M-wave and an almost flat line in electromyographic activity) and still being comfortable for the participant. This methodology of electrode placement was replicated for each muscle. Figure 1 gives an overview of all electrode locations and their associated EMG recordings.

**Figure 1:**
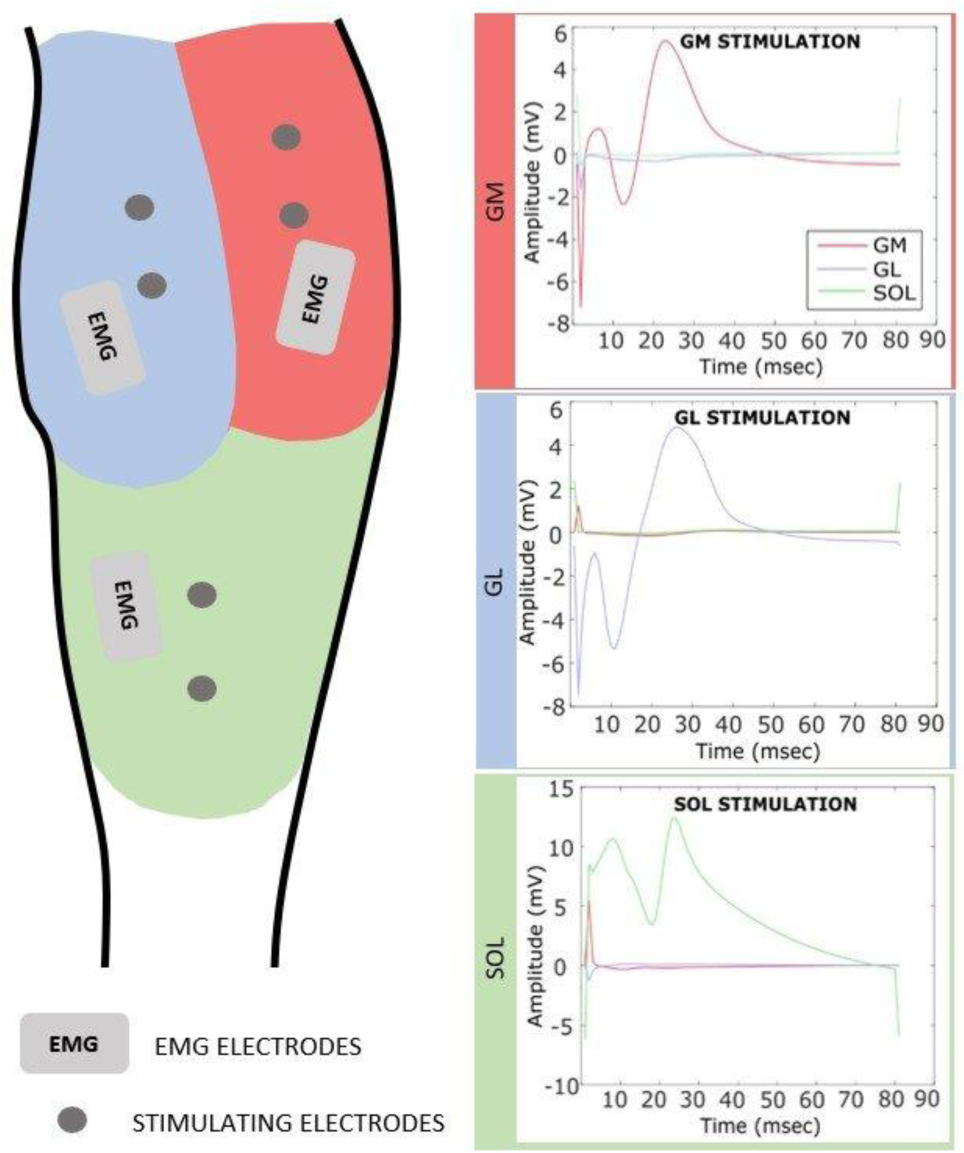
Schematic representation of the location of the EMG recording and stimulating electrodes on gastrocnemius medialis (GM; red), gastrocnemius lateralis (GL; blue) and soleus (SOL; green). The left panel shows the electrodes positioning. The right panel shows the electromyographic activity during individual stimulation of GM, GL and SOL, resulting in an M-wave in the targeted muscle without contribution from the other two muscles, visually checked by an almost flat line.

Each muscle received 3 trains of 30 pulses of 50 µs each, frequency 25 Hz, one in each foot position [toes-neutral, toes-in and toes-out], with at least a two-minute rest interval between each stimulation trial. For the toes-neutral, the footplate was aligned in the sagittal plane. For the toes-in and toes-out foot positions, the footplate was rotated inwards and outwards by 30°, respectively. During every stimulation, torque, ultrasound imaging and EMG activity were recorded. The order of muscles stimulated and foot positions were randomized.

2D-ultrasound images were recorded in the sagittal plane with the ultrasound probe (referenced above), placed distally on the AT, the calcaneal notch being used as a reference landmark for probe position. Probe orientation was consistently determined by ensuring the tendon borders were parallel on the ultrasound image. A piece of hypoechoic tape was also attached to the skin at the level of the calcaneal notch to correct for any possible displacement of the probe over the skin. Ultrasound radio frequency (RF) data were collected at 70 frames/s using the Telemed Ultrasound Research Interface (ArtUs RF-Data Control, v1.4.4).

### 2.3. Data processing

#### 2.3.1. **Torque**

Torque was analyzed using Matlab 2020b (MathWorks Inc., Natick, MA) and low-pass filtered at 10Hz. For the MVC’s, maximal torque was calculated as the highest value over a 500ms moving average window. For the stimulation trials, the average torque during stimulation was calculated (except for the first and last 0.20s).

#### 2.3.2. Speckle tracking

2D ultrasound images were analyzed using a validated ultrasound speckle tracking algorithm (MathWorks Inc., Natick, MA) (Fig 2) (Slane and Thelen 2014, Dandois et al. 2021). Superficial, middle and deep layers of the AT were tracked by computing the absolute displacement from RF data (upsampled by a factor of 2 and 4 in the along-fiber (x) and transverse (y) directions) (Parker et al. 1983) to increase spatial density of correlation functions (Konofagou and Ophir 1998). A manually selected region of interest (ROI) with 3 rows (10mm length) and 13 columns of kernels was determined. Frame-to-frame displacements during the stimulation period were computed using a 2D normalized cross-correlation technique. RF data within a kernel centered at the current node location were cross-correlated with RF data in a search window at the same location in the subsequent frame. A correlation threshold of r = 0.7 was used to discriminate valid displacement information (Farron et al. 2009, Korstanje et al. 2010). Kernel size, search window sizes and cross-correlation threshold were based on previous publication (Stenroth et al. 2019) and adjusted if tracking failed. Cumulative 2D displacements for each node were obtained during the stimulation period and tracking was repeated in the reverse direction as well. The weighted average of forward and backward tracking provided the nodal displacement trajectories and so the absolute displacement of the superficial, middle and deep AT layers. The mean of the absolute displacements of all three sublayers was also calculated. Intra-tendinous sliding was calculated as the differential displacement between the superficial and deep layers. Normalized displacement values for all AT layers were calculated as the relative contribution of the AT sublayer displacement to the sum displacement of all AT layers together. ICC values for within- and between-session reliability for intra-tendinous sliding were respectively 0.908 (95% CI: 0.815 – 0.954) and 0.742 (95% CI: 0.405 – 0.852). For mean absolute AT displacement, these were respectively 0.968 (95% CI: 0.936 – 0.984) and 0.755 (95% CI: 0.509 – 0.877), indicating overall a good to excellent reliability. More detailed information about reliability analysis can be found in supplementary material.

**Fig 2:**
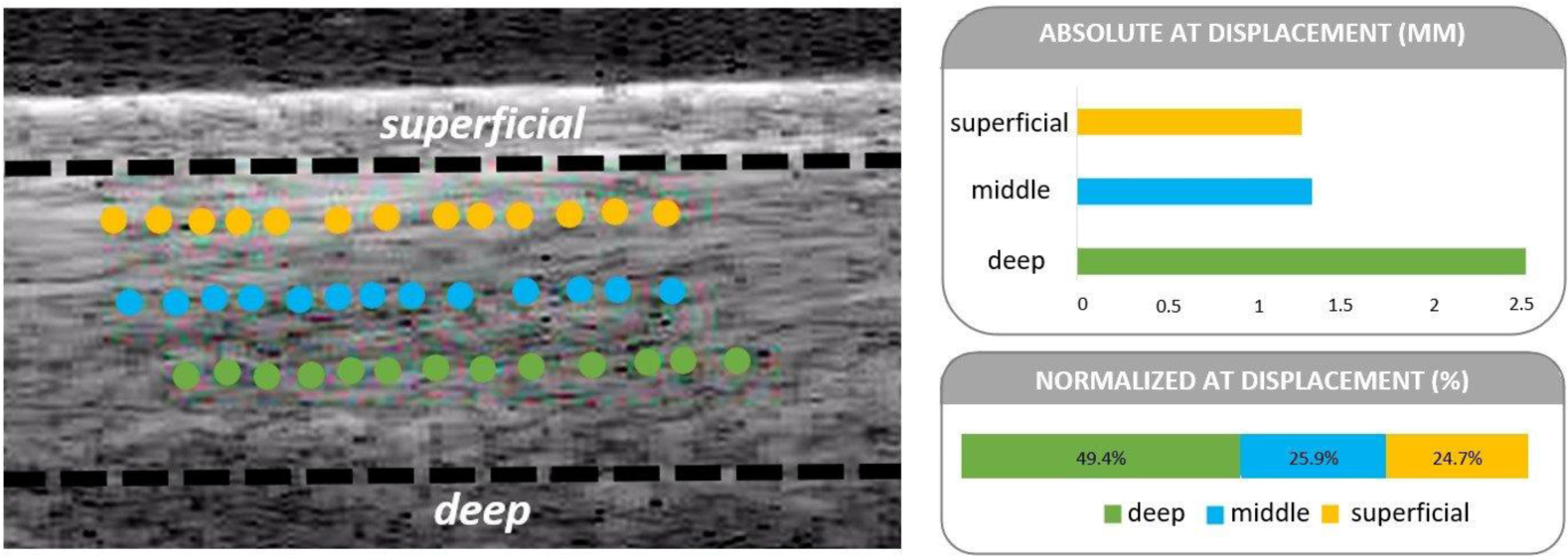
Ultrasound image at the end of a muscle stimulation. A speckle tracking algorithm is used to track the absolute displacement of the superficial (yellow), middle (blue) and deep (green) AT sublayer, each layer consisting of 13 kernels. Dotted black lines represent the superficial and deep AT borders. AT normalized displacement is calculated as the relative contribution of the AT sublayer displacement to the sum displacement of all AT layers together. Intra-tendinous sliding is calculated as the difference between the absolute displacement of the superficial and deep layer.

#### 2.3.3. AT twist estimation based on speckle tracking

Based on the twist classifications of Edama et al. (2015) or Pekala et al. (2017), our subjects were divided into the different twist categories from the normalized displacement values of AT layers during stimulation of the three muscles in neutral foot position. Criteria were first based on the normalized displacement of the AT deep layer followed by the displacement of the AT superficial layer.

##### Classification based on Edama’s twist definition

Based on Edama and collegeaus, SOL inserts into the deep layer for Type I twist, whereas the deep layer is both GL and SOL for Type II; and is only GL for Type III (Fig 3). Based on this, if SOL-stimulation induced the highest normalized deep displacement compared to GM or GL stimulation (Δ > 5%), the subject was classified as Type I. If GL stimulation induced the highest deep displacement (Δ > 5%), it was Type III. If GL and SOL stimulations showed very little difference (Δ < 5%), it was Type II.

**Figure 3:**
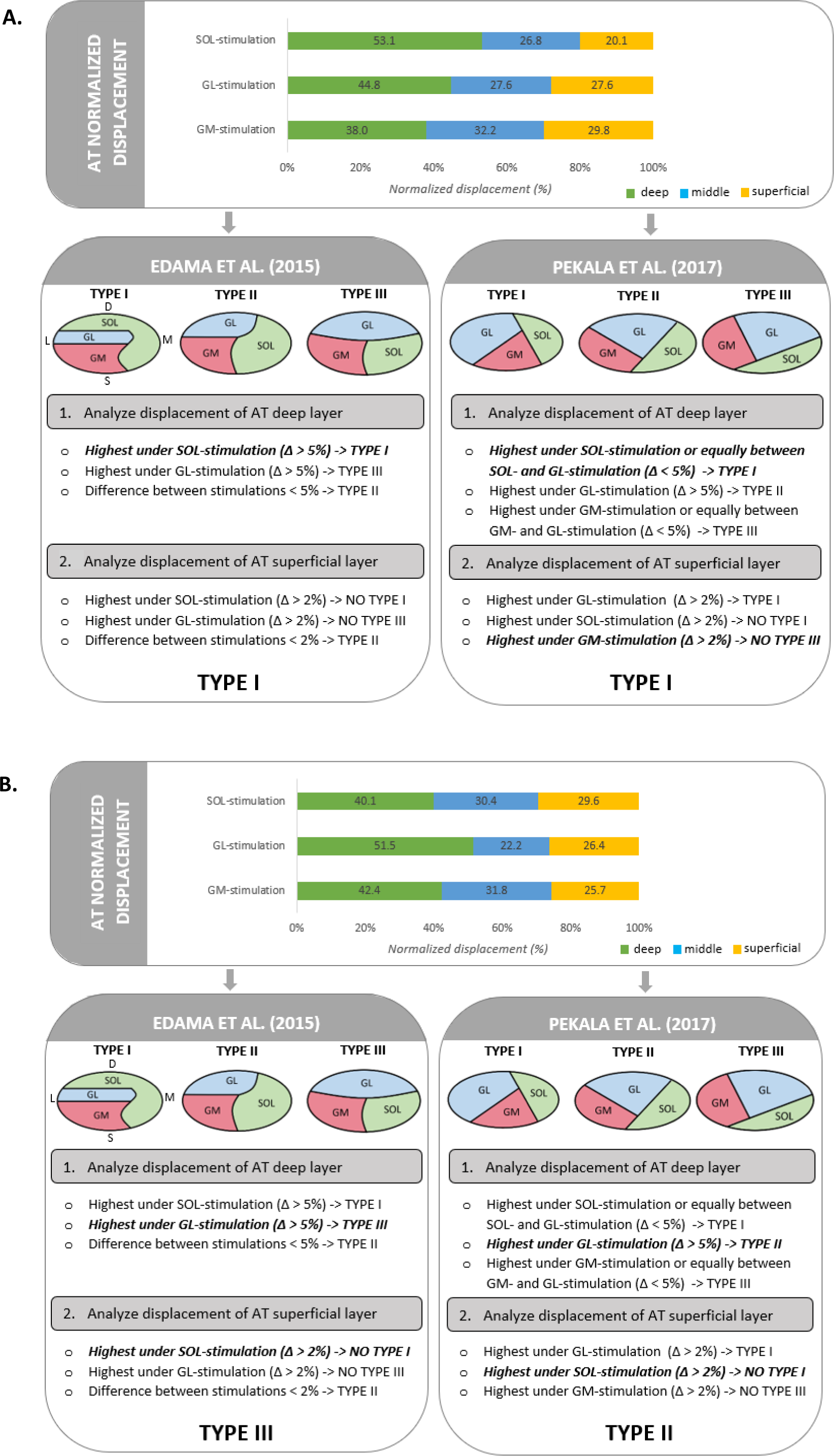
Example of two datasets to classify the tendon twist following the different criteria, based on the two twist classifications proposed by Edama et al. (2015) and Pekala et al. (2017). The graph on top represents the normalized AT displacement of the deep, middle and superficial layer of one representative subject. For each twist classification, a schematic representation of all subtendons (GM = gastrocnemius medialis, GL = gastrocnemius lateralis, SOL = soleus) at the insertion into the calcaneal bone is shown: D represents the deep part, S represents the superficial part, L represents the lateral part and M represents the medial part of the Achilles tendon. Criteria were first based on the displacement of the AT deep layer followed by the displacement of the superficial layer. Criteria that corresponded correctly are indicated in bold. A) Subject classified as Type I in both classifications. B) Subject classified as Type III according to Edama et al. (2015) and Type II according to Pekala et al. (2017).

Then, the superficial layer was used to confirm or infirm the first classification. Based on Edama and colleagues, the superficial part is never really occupied by only one subtendon, but SOL cannot be seen in the superficial layer for Type I twist, it can be either GM and SOL for Type II, and it cannot be GL for Type III. Hence, if SOL stimulation induced the highest normalized superficial displacement, Type I was eliminated. If the highest displacement occurred under GL stimulation, Type III was eliminated. If GM and SOL stimulations showed very little difference in normalized superficial displacement (Δ < 2%), it was categorized as Type II.

##### Classification based on Pekala’s twist definition

The same method was repeated following Pekala’s classification. Based on Pekala and colleagues, GL and SOL insert into the deep layer for Type I twist, whereas the deep layer is dominated by GL for Type II; and is comprised of GM and GL for Type III. Based on this, if SOL-stimulation induced the highest normalized deep displacement (Δ > 5%) compared to GM and GL stimulation, or if normalized deep displacement was equal between GL and SOL- stimulation (Δ < 5%), the subject was classified as Type I. If GL stimulation induced the highest deep displacement (Δ > 5%), it was categorized as Type II. If normalized deep displacement showed little difference between GM and GL stimulation (Δ < 5%) or was highest under GM stimulation (Δ > 5%), it was categorized as Type III.

Then, the superficial layer was used to confirm or infirm the first classification. Based on Pekala and colleagues, Type I is the only twist type where GL contributes to the superficial layer, the smallest contribution of GM in the superficial layer is in Type III and the smallest contribution of SOL in the superficial layer is in Type I. Hence, if GL stimulation induced the highest normalized superficial displacement, this was classified as Type I. If the highest displacement occurred under SOL stimulation, Type I was excluded. If the highest displacement occurred under GM stimulation, Type III was excluded.

#### 2.3.4. **EMG**

EMG signals were band-pass filtered (20–500 Hz), full-wave rectified and analyzed during the first 80ms of the stimulation pulse train. All M-waves were visually checked and trials identified as lower quality were removed. As a result, 2 participants were excluded from further analysis. Among these 2 subjects, one stimulating condition was identified as being of lower quality; therefore, all data from these subjects were removed. The M-wave latency was manually determined by using the peak-to-peak function where the top and bottom of the M-wave were indicated. This enabled us to compute the amplitude of the M-wave, as well as the signal amplitudes of the other muscles, which was expected to be close to zero. ICC values for within- and between-session reliability for M-wave were respectively 0.99 (95% CI: 0.98 – 1) and 0.68 (95% CI: 0.40 – 0.87) (see supplementary material).

### 2.4. Statistics

One-way repeated measures ANOVA was used to investigate the effect of the stimulated muscle [GM, GL and SOL] on mean absolute AT displacement, normalized AT layers absolute displacement, intra-tendinous sliding, torque and M-wave amplitude. Similarly, one-way repeated measures ANOVA was used to investigate the effect of the horizontal foot position [toes-neutral, toes-in and toes-out] on these parameters. Where appropriate, post-hoc analyses were performed using the Bonferroni test. The level of significance was set at p ≤ 0.05. Data are presented as mean ± standard deviation.

## RESULTS

### EFFECT OF MUSCLE DURING NEUTRAL STIMULATIONS

There was a main effect of muscle on the **absolute AT displacements** (F(2,56) = 3.485, p < 0.001), with absolute displacements being lower under GL stimulation compared to GM and SOL stimulation (both p < 0.001), and no difference between GM and SOL stimulations (p = 1.000). Interestingly, regardless of the muscle stimulated, the same pattern of relative displacement between tendon layers occurred with the deep layer moving more than the middle and superficial layers (all p < 0.007) (Fig 4). When expressed in **normalized displacement values**, i.e. each layer expressed as a percentage of the sum of the displacement of the three layers, no differences were found between muscle stimulations (all p > 0.134). Finally, **intra- tendinous sliding**, calculated as the relative displacement between the deep and superficial layer, revealed a main effect of muscle (F(2,56) = 7.386, p = 0.037). Intra-tendinous sliding was lower with GL stimulation compared to GM stimulation (p = 0.003) and SOL stimulation (p = 0.005), but no difference was found between GM and SOL stimulation (p = 1.000). Of note, there was a main effect of muscle on the **torque** during stimulation (F(2,52) = 9.736, p < 0.001), with torque being lower during GL stimulation than during GM and SOL stimulation (p = 0.024 and p = 0.002 respectively). No differences were found between the torques during GM and SOL stimulation (p = 0.149).

**Figure 4:**
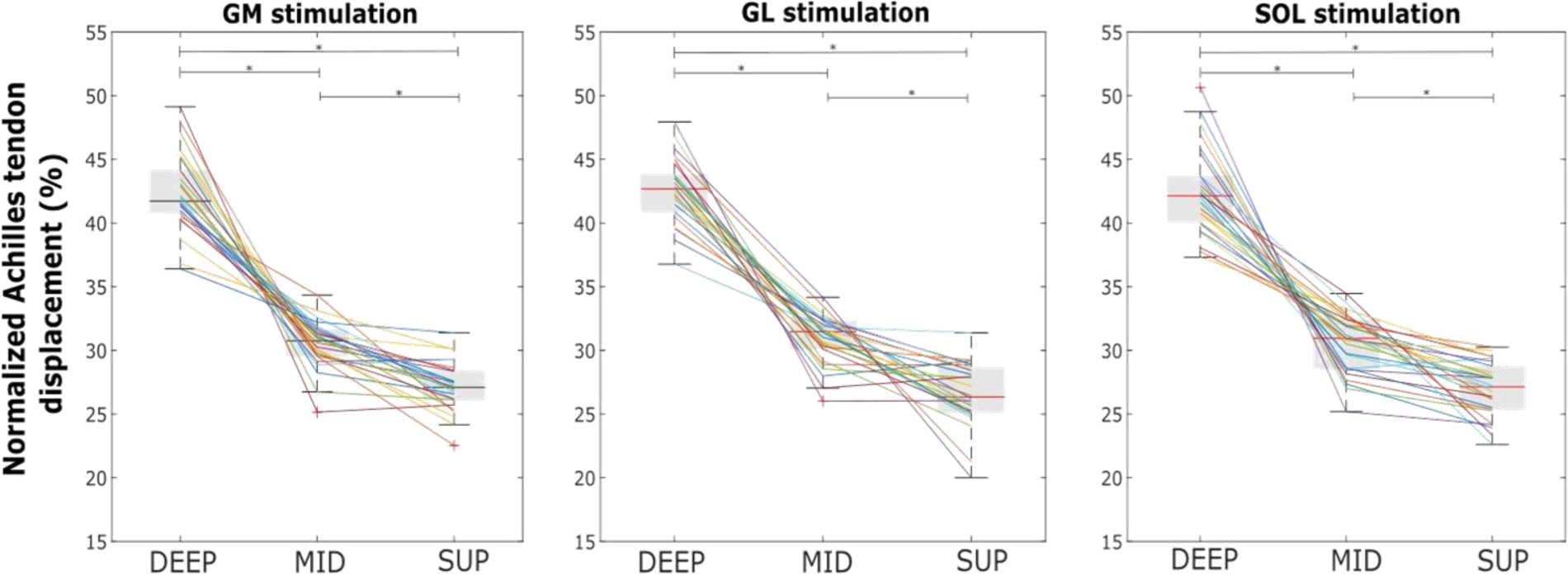
Normalized AT absolute displacements of the deep (DEEP), middle (MID) and superficial (SUP) layer during stimulations of GM, GL and SOL with the foot in neutral position. Each colored line represents one subject. *: p < 0.001. Deep, mid and up refer to the three layers of the Achilles tendon that were tracked with a speckle tracking analysis on the ultrasound images.

### EFFECT OF FOOT POSITION DURING STIMULATIONS

There was a main effect of horizontal foot position on the **absolute AT displacements** (F(2.58) = 5.510, p = 0.006), which was higher with toes-out (1.15 ± 0.68 mm) compared to toes-in (0.99 ± 0.52 mm, p = 0.021). No differences were found between toes-neutral (1.02 ± 0.62 mm) and the two other foot positions (both p > 0.081). A main effect of foot position was also found for **intra-tendinous sliding** (F(2,58) = 4.027, p = 0.023). Intra-tendinous sliding was higher in toes-out (0.48 ± 0.23 mm) compared to toes-neutral (0.41 ± 0.22 mm, p = 0.003). No differences were found between toes-in (0.42 ± 0.21 mm) and the two other foot positions (both p > 0.091). Figure 5 shows the effect of horizontal foot position on absolute AT displacements and intra- tendinous sliding. Notably, there was a main effect of foot position on **M-wave amplitude** regardless of the muscle (F(2,178) = 5.235, p = 0.006), which was higher with toes-out (3.01 ± 2.26 mV) compared to toes-neutral (2.82 ± 2.12 mV, p = 0.003). No differences were found between toes-in (2.83 ± 2.14 mV) and the two other foot positions (both p > 0.098).

**Figure 5:**
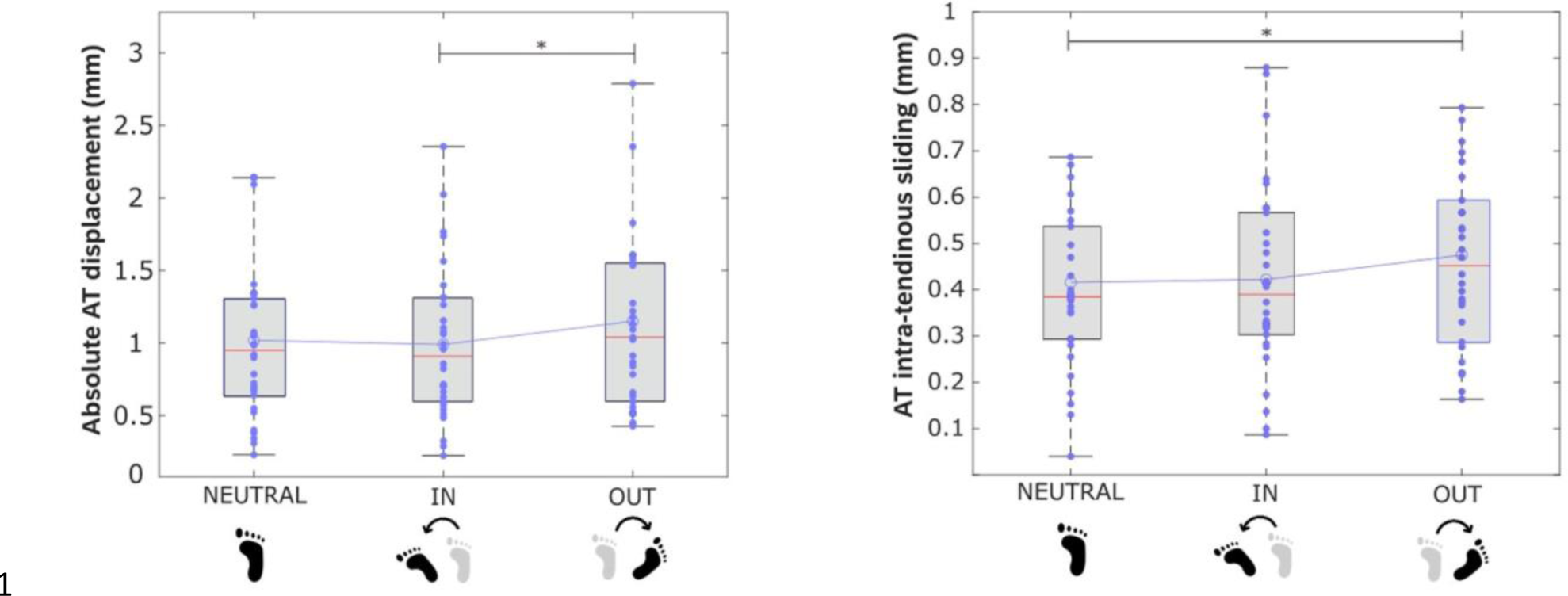
Boxplots showing the effect of horizontal foot position [toes-neutral (NEUTRAL), toes-in (IN) and toes-out (OUT)] on absolute AT displacement (left) and intra-tendinous sliding (right). Every dot represents the average value of GM, GL and SOL stimulation in that specific foot position for one subject. The blue line connects the means with each other, the red line represents the median. * represents statistical difference between foot positions (p < 0.05).

### AT TWIST ESTIMATION

Two twist type distributions were generated based on Edama’s and Pekala’s classifications (Edama et al. 2015, Pekala et al. 2017). For Edama’s classification, 22 of 30 subjects met our definition criteria for both deep and superficial AT layer displacement. For Pekala’s classification, this was 21 subjects. The detailed classifications are shown in the table below.

Bringing both classifications together, we were able to make a distinction between subjects with a low amount of twist (Type I and/or Type II in both classifications) and subjects with a high amount of twist (Type II and/or Type III in both classifications). This resulted in a “low twist” group of 19 subjects and a ”high twist” group of 11.

## DISCUSSION

In this study, we investigated the AT displacement patterns during selective stimulation of the triceps surae muscles across various horizontal foot positions to enhance our understanding of the AT substructure. As hypothesized, we found that SOL stimulation yielded the highest absolute displacement of all AT sublayers, followed by GM stimulation and GL stimulation. Regarding the horizontal foot position, the external foot position resulted in higher absolute AT displacement patterns and also higher intra-tendinous sliding. Interestingly, we were able to classify the participants into two groups, i.e. with “low” or “high” twist based on AT ultrasound recordings. This classification followed the twist description given by Edama et al. (2015) and Pekala et al. (2017).

During SOL stimulation, we found the largest absolute displacement and the largest torque. About the last mentioned, torque during SOL stimulation was the highest followed by GM and GL, in agreement with Khair et al. (2022). This is to be expected, as SOL has the largest physiological cross-sectional area, which is closely correlated with muscle force generation (Fukunaga et al. 1997). The same trend was found for the AT absolute displacement. Lehr et al. (2021) found that absolute displacement was higher during SOL stimulation compared to GM stimulation, and Khair et al. (2022) found that GM produced higher absolute displacements compared to GL stimulation. Our study, which included independent stimulation of all three muscles, showed a similar pattern, with SOL stimulation generating the highest absolute displacement, followed by GM stimulation and the lowest during GL stimulation. Interestingly, irrespective of the muscle stimulated, and consistently with the afore-mentioned studies, the deep AT layer showed more displacement than the superficial layer. Furthermore, we observed relative sliding between tendon layers, which were already known to occur in contexts beyond individual muscle stimulation, such as passive motions, or voluntary (eccentric) contractions (Arndt et al. 2011, Slane and Thelen 2014, Franz et al. 2015, Bogaerts et al. 2017). More specifically, we found that intra-tendinous sliding was highest under SOL stimulation and lowest under GL. Lehr et al. (2021) found 49% more intra-tendinous sliding during SOL compared to GM stimulation with the foot in 20 degrees of plantarflexion. This ankle position probably allows for larger subtendon tissue displacements and reduced passive tension due to the AT being slack. Their results (see Figure 5) suggest a smaller difference when the foot was neutral, aligning more closely with the 6.5% difference we found between SOL and GM stimulation. Khair et al. (2022) did find similar sliding between GM and GL stimulation, i.e. only a difference of 4.5%, whereas we found a difference of 43.7% in intra-tendinous sliding between these muscle stimulations. However, they stimulated at lower intensities. It’s plausible that significant differences in intra-tendinous sliding may emerge only at higher intensities, potentially exceeding a specific threshold of force transmission.

Our data revealed important inter-individual differences when normalizing AT sublayers to overall displacement. We found similar patterns for the normalized displacement of all three layers between muscle stimulations; the deep layer moving relatively the most followed by the middle and then superficial layer. However, caution is warranted in interpreting these average values, as individual differences become apparent upon closer inspection. In 3 of 30 subjects, normalized deep displacement was substantially higher during GM stimulation compared to the normalized deep displacement during the other two muscle stimulations, in 18 subjects this was highest during GL stimulation, and in the remaining 11 subjects during SOL stimulation. Additionally, in half of the subjects, the superficial layer moved more than the middle layer during specific stimulations. This highlights the importance of interpreting such data on an individual basis and not as averages only, as there are many inter-individual differences that could potentially be associated with specific AT twisting types.

The external horizontal foot position resulted in higher absolute displacement and intra- tendinous sliding. No differences in torque were observed during stimulation between foot positions, indicating this is not a critical parameter influencing AT behaviour across foot positions. Hence, these variations in AT behaviour are rather associated with the intrinsic properties of the AT itself. It could be that the horizontal foot position influences the amount of slack in the muscle-tendon unit: greater is the slack, lesser is the force transmission to the bone. Outward foot rotation counteracts the natural AT twist, potentially unwinding it and releasing the tight connections between the fascicles, facilitating sliding between them. Additionally, it has been shown that subtendons not only twist around each other but also exhibit individual self-twisting (Pekala et al. 2017). This intrinsic twist within each subtendon may be influenced by the horizontal foot position, potentially impacting the overall stiffness of the AT. Our findings align with literature showing increased intra-tendinous sliding during isometric plantarflexion in healthy individuals and those with Achilles tendinopathy, though the causative factor remained unknown (Crouzier et al. 2022, Lecompte et al. 2024). To gain a more comprehensive understanding, the impact of external foot positioning on AT displacement should be explored in dynamic voluntary, where more varied triceps surae behaviour could contribute to further insights into intra-tendinous sliding.

The current study is the first to try to make an in vivo AT twist classification based on ultrasound AT displacement data. We were able to identify people with a low and a high twisted AT, a notable achievement given the difficulty of measuring this feature. Recently, Cone et al. (2023) pioneered the investigation of AT twist using high-field 7-Tesla MRI in 10 healthy young subjects where they achieved to reconstruct the distal 50mm of each subtendon. While their method is costly and time-consuming, our study, though capturing a smaller part of the distal AT, introduces a novel exploratory in-vivo method to estimate AT twist. Research efforts, as demonstrated by Cone et al. (2023) and our study, are crucial for enhancing our understanding of the relationship between specific AT morphology, behavior, and injury mechanisms in-vivo. To date, only cadaveric and simulation studies have previously attempted to answer the question of how the twist influences AT behaviour, recognizing its potential role in Achilles tendinopathy onset (Bojsen- Møller et al. 1985, Lyman et al. 2004, Lersh et al. 2012, Edama et al. 2017). However, the precise biomechanical effects of AT twist and whether high or low twist is linked to Achilles tendinopathy remain unclear. Finite element modelling was used to investigate the influence of twist on AT behaviour. Studies incorporating greater geometric twist in finite element models suggest a redistribution of internal stresses, enhancing the AT’s capacity to withstand greater loads (Shim et al. 2018, Knaus et al. 2021). Funaro et al. (2022) developed subject-specific 3D models for dynamic rehabilitation exercises, offering a functional perspective of the twist in the context of Achilles tendinopathy. They found that the least twisted geometry exhibited the highest peak strain, potentially increasing the risk of injury. Conversely, some studies suggest that a higher twist may increase the risk of Achilles tendinopathy due to increased compression of the vascular supply to the tendon or increased internal compression (Pekala et al. 2017, Edama et al. 2021, Pringels et al. 2023). Handsfield et al. (2020) suggested an optimum twist, neither excessively high nor low, for optimizing rupture load and stress distributions. These conflicting findings underscore the necessity for further in-vivo investigations to comprehensively explore the relationship between twisted morphology and muscle-tendon behaviour.

This study has several limitations. First, a 2D speckle tracking algorithm was employed to investigate the 3D structure of the AT, potentially resulting in an under- or overestimation of AT displacement. Additionally, conducting only one trial per condition may increase the risk of measurement error in AT displacement. However, the within-session reliability was good for AT displacement and all subjects that were measured twice (n = 4) were consistently classified in the same twist pattern group for both sessions, confirming the method’s reliability. Next, we used an indirect method to investigate the AT twist, which may not be optimal. The electrode location and the stimulation intensity were determined to generate a maximal M-wave in the neutral position. In the external foot position, we observed an increase in M-wave amplitude. Hence, it is possible that the external foot position had a small impact on the nerve position, relative to the stimulating electrode position or a small impact in the recording volume.

## CONCLUSION

This study is the first that stimulated all triceps surae muscles individually, while simultaneously recording ultrasound images of the distal AT. Based on the displacement of the AT sublayers, we were able to classify subjects as having either a *low* or *high* AT twist. As this is very exploratory and cannot be easily validated, further research is needed to understand the complexity of the 3D twisted structure of the AT in-vivo. This will contribute to a more comprehensive understanding of its impact on AT properties and behaviour.

## ACKNOWLEDGEMENTS

We would like to thank Madeline Ella Ford for her assistance during the measurements. The author declares no conflict of interest.

## Data Availability Statement

The data that support the findings of this study are available from the corresponding author upon reasonable request.

## AUTHOR CONTRIBUTIONS

<u>Laura Lecompte)</u> Concept/design, pilot testing, acquisition of data, data analysis/interpretation, figures and graphs, drafting and writing of the manuscript <u>Marion Crouzier)</u> concept/design, data analysis/interpretation, critical revision of the manuscript, approval of the manuscript <u>Stéphane Baudry)</u> help during pilot testing with his expertise, critical revision of the manuscript, approval of the manuscript <u>Benedicte Vanwanseele)</u> concept/design, data analysis/interpretation, critical revision of the manuscript, approval of the manuscript

## Notes

### Competing Interest Statement

The authors have declared no competing interest.

